# Using *Vibrio natriegens* for high-yield production of challenging expression targets and for protein deuteration

**DOI:** 10.1101/2023.11.03.565449

**Authors:** Natalia Mojica, Flore Kersten, Mateu Montserrat-Canals, G. Robb Huhn, Abelone M. Tislevoll, Gabriele Cordara, Ken Teter, Ute Krengel

## Abstract

Production of soluble proteins is essential for structure/function studies, however, this usually requires milligram amounts of protein, which can be difficult to obtain with traditional expression systems. Recently, the Gram-negative bacterium *Vibrio natriegens* appeared as a novel and alternative host platform for production of proteins in high yields. Here, we used a commercial strain derived from *V. natriegens* (Vmax^TM^ X2) to produce soluble bacterial and fungal proteins in milligram scale, which we struggled to achieve in *Escherichia coli*. These proteins include the cholera toxin (CT) and *N*-acetyl glucosamine binding protein A (GbpA) from *Vibrio cholerae*, the heat-labile enterotoxin (LT) from *E. coli* and the fungal nematotoxin CCTX2 from *Coprinopsis cinerea*. CT, GbpA and LT are secreted by the Type II secretion system in their natural hosts. When these three proteins were produced in Vmax, they were also secreted, and could be recovered from the growth media. This simplified the downstream purification procedure and resulted in considerably higher protein yields compared to production in *E. coli* (6– to 26-fold increase). We also tested Vmax for protein deuteration using deuterated minimal media with deuterium oxide as solvent, and achieved a 3-fold increase in yield compared to the equivalent protocol in *E. coli*. This is good news since isotopic labeling is expensive and often ineffective, but represents a necessary prerequisite for some structural techniques. Thus, Vmax represents a promising host for production of challenging expression targets and for protein deuteration in amounts suitable for structural biology studies.

## INTRODUCTION

Obtaining structural information on toxins and other virulence factors from bacteria and fungi is essential to understand the molecular mechanisms behind infections. Structural biology studies often rely on the production of the target protein in high amounts (milligram scale), which is usually achieved by expression in *Escherichia coli.*^1^ This Gram-negative bacterium is the first choice for protein production in many laboratories due to its fast growth in inexpensive media, its well-characterized genome, and the availability of a wide range of commercial *E. coli* strains for different applications.^2^ However, expression yields are not always sufficient; in addition other challenges abound, such as generation of insoluble aggregates.^3^ *E. coli* strains can also be used for the production of deuterated proteins, which is highly useful for techniques involving neutron scattering, such as neutron crystallography and small-angle neutron scattering (SANS).^4^ In deuterated proteins, a significant number of the hydrogen atoms are substituted for deuterium. Deuterated reagents are expensive, highlighting the importance of an expression system that can efficiently produce high yields of protein. Overcoming challenges related to protein production often requires significant optimization of the growth conditions, modification of the constructs or the use of different bacterial strains, which is a time-consuming process and not always successful.

As an alternative to *E. coli*, a new potential expression host has emerged: *Vibrio natriegens.*^5^ This Gram-negative marine bacterium can grow twice as fast as *E. coli* under optimal conditions.^6^ Such rapid growth requires a very efficient protein synthesis machinery, which has been mainly attributed to the higher number of ribosomes produced by *V. natriegens* in the exponential phase – 115 000 versus 70 000 in *E. coli.*^7^ In addition, *V. natriegens* has its genome distributed between two chromosomes, thus replication can occur in parallel from two origins of replication.^8^ Its fast growth and ability to rapidly synthesize proteins makes *V. natriegens* an interesting species for protein production. In 2017, an engineered strain known as Vmax™ X2 (originally Vmax™ Express) was made commercially available for the expression of genes under the control of an arabinose or isopropyl β-D-1-thiogalactopyranoside (IPTG) inducible T7 promoter, allowing the use of *E. coli*-based plasmids in *V. natriegens.*^5^ Since then, different proteins have been produced using Vmax™ X2, including membrane proteins from *Vibrio cholerae,*^9^ recombinant enzymes cloned into pET vectors^10^ and an insect metalloproteinase inhibitor fused to *N*-acetylglucosamine-binding protein A (GbpA),^11^ which was used as secretion tag. To the best of our knowledge, *V. natriegens* has not yet been used for protein perdeuteration.

In our own work, we have experienced on a number of occasions the limitations of *E. coli* for producing soluble, active proteins in sufficient amounts for structural studies. One problematic target is the cholera toxin (CT), the main virulence factor of *V. cholerae*. CT is composed of a catalytic A subunit (CTA) and a homopentamer of cell-binding B subunits (CTB) that are assembled to an AB_5_ toxin in the periplasm. Thereafter, the multimeric toxin is secreted into the extracellular environment by the *V. cholerae* type II secretion system (T2SS).^12–14^ When produced in *E. coli*, we could not recover CT from the growth medium. Only low yields of toxin (0.2 mg per L culture) were obtained from the periplasmic space, in a mixture of intact holotoxin and free CTB. We reasoned that recombinant CT would be secreted by the *V. natriegens* T2SS,^11^ which could improve holotoxin yield and purity. *V. natriegens* therefore appeared like a suitable system to produce CT. In addition, we tested production of two other proteins secreted by the T2SS: GbpA from *V. cholerae* and heat-labile enterotoxin (LT) from enterotoxigenic *E*. *coli* (ETEC). We also tested *V. natriegens*—with success—for protein deuteration, which produced increased yields when compared to *E. coli* expression systems, opening the door to neutron-based techniques^4^. In a final test, we extended the work to non-secreted proteins, expressing a fungal nematotoxin from *Coprinopsis cinerea* (CCTX2).^15^ When compared to expression in *E. coli*, Vmax™ X2 significantly improved the yields of all tested proteins and in some cases considerably simplified the purification process. *V. natriegens* thus represents a promising system for the production of challenging protein targets.

## MATERIALS AND METHODS

### Constructs and bacterial strains

Plasmids encoding CT, LT, GbpA and *C. cinerea Toxin 2* (CCTX2) were used for protein production in both *E. coli* and Vmax™ X2. N-terminal signal sequences directed synthesized CT, LT and GbpA to the periplasmic space, where they were cleaved from the mature proteins, whereas CCTX2 remains in the cytoplasm of the bacterial expression hosts.

Professor Randall K. Holmes kindly provided us with a pARCT5 vector (derived from the pAR3 vector)^16^ encoding the *ctxAB* operon under the control of an L-arabinose-inducible promoter (Figure S1A). The translated genes contained the signal sequence from *E. coli* LT-IIB. This tag directs CTA and CTB to the periplasmic space more efficiently than their native CT secretion signals.^17^

The operon encoding LT from ETEC strains of porcine origin (also known as pLT) was synthesized by Genscript® with LT-IIB signal sequences, provided in a pUC57 vector, and subcloned into a mutated version of pARCT5 containing only two (instead of three) NcoI restriction sites. These restriction sites were used to replace the DNA coding for CTA and CTB with the coding sequences for LTA and LTB (*i.e.*, *eltA* and *eltB*), respectively (Figure S1B).

The gene encoding GbpA (Uniprot ID: Q9KLD5, residues 24-485) was codon-optimized for *E. coli* and cloned into pET26b(+) by GenScript® (Leiden, Netherlands) using restriction sites NcoI and XhoI. The natural signal peptide of the protein was substituted by the pelB leader sequence present in the vector, which in *E. coli* signals the protein for periplasmic localization but in Vmax™ X2 directs the secretion of the protein into the culture media (Figure S1C). The vector originally contained a C-terminal His_6_-tag, but introduction of a stop codon at the end of the insert prevented expression of the tag.

For CCTX2, vector pET24b(+), which contains the wild-type *cctx2*-gene under control of an IPTG-inducible T7 promoter (Figure S1D), was provided by Professor Markus Künzler (ETH Zurich).

### Transformation

All plasmids were introduced into chemically competent Vmax™ X2 cells (TelesisBio, San Diego) by heat shock, following the manufacturer’s guidelines. Briefly, 100-200 ng of plasmid DNA were incubated with Vmax™ X2 competent cells on ice for 30 min before heat shock at 42°C for 45 seconds in a water bath. The cells were immediately returned to ice for 2-5 min and then transferred to a pre-warmed 14 mL Falcon® tube containing 950 μL of Vmax™ X2 chemicompetent cell recovery medium and incubated at 30°C for 2 h in a shaker. Cells were plated on Luria Bertani (LB) agar plates containing the appropriate antibiotics (12.5 μg/mL chloramphenicol (CAM) for cells transformed vectors encoding CT and LT, or 100 μg/mL kanamycin (Kan) for GbpA and CCTX2-encoding vectors) and incubated overnight at 30°C. Single colonies were taken to grow overnight cultures in LB+v2 medium (LB media supplemented with 204 mM NaCl, 4.2 mM KCl, and 23.14 mM MgCl_2_ (v2 salts))^18^ with corresponding antibiotics. Subsequently, we prepared glycerol stocks that were stored at – 80°C.

### Recombinant expression in E. coli

*Expression of ctxAB and eltAB.* Expression was essentially performed as described previously.^19^ Briefly, OverExpress™ C43 (DE3) cells (Sigma) harboring pARCT5 or pARpLT5 were grown overnight at 30°C in Terrific Broth (TB) medium containing 25 μg/mL CAM. Cultures were diluted 1/50 in TB medium, grown until the optical density at 600 nm (OD_600_) reached 2.0 and induced with 0.2% L-arabinose (Sigma) at 37°C. After 3 h, the cells were harvested by centrifugation (6,000 x *g*, 20 min, 4 °C) and resuspended in 1/40^th^ volume of Talon A buffer (50 mM sodium phosphate pH 8.0, 300 mM NaCl), supplemented with c*O*mplete™ protease inhibitor cocktail (Roche), 1 mg/mL polymyxin B sulfate (Sigma) and benzonase (Sigma). This solution was incubated at 37°C for 15 min with shaking followed by centrifugation (8,000 x *g*, 20 min, 4°C). The supernatant (containing the periplasmic extract) was filtered through a 0.22 μm filter (polyethersulfone (PES) membrane, VWR) and used immediately for further purifications steps.

*Expression of gbpA*. GbpA production was carried out essentially as described by Sørensen *et al*.^20^ BL21 Star (DE3) cells transformed with the codon-optimized GbpA-encoding pET26-b(+) vector were grown for 6 h at 37°C, 220 rpm, in 2.5 mL LB medium containing 50 μg/mL Kan. 200 μL of the pre-culture was diluted in 25 mL (1/125 dilution) minimal medium M9glyc+^21^ (see Table S1 for recipe) and grown for 14 h at 37°C, 130 rpm. The main culture was started by adding 225 mL of M9glyc+ medium to the culture and adjusting the concentration of antibiotic back to 50 μg/mL Kan. When the OD_600_ reached 2 to 3, IPTG was added to a final concentration of 1 mM to induce expression. After incubation for 18 h at 20°C, 130 rpm, cells were harvested by centrifugation at 10,000 *x g* for 30 min at 4°C. GbpA was isolated from the periplasmic space by osmotic shock. Briefly, the pellet was resuspended in 5 mL sucrose solution (25% w/v sucrose, 20 mM Tris-HCl pH 8.0, 5 mM ethylenediaminetetraacetic acid (EDTA)) per gram cells. The cell suspension was incubated on ice for 30 min under stirring, followed by centrifugation at 10,000 *x g* for 30 min at 4°C. The supernatant was saved as sucrose fraction for further processing. The pellet was resuspended in 5 mL per gram of cells of a hypotonic solution (5 mM MgCl_2_, 0.25 mg/mL lysozyme from chicken egg white >40,000 units/mg protein (Sigma) and 1 mM phenylmethylsulfonyl fluoride). The suspension was again incubated on ice for 30 min under stirring, followed by centrifugation at 10,000 *x g* for 30 min at 4°C. The supernatant was pooled together with the sucrose fraction and filtered through a 0.22 μm PES membrane (VWR) and used immediately or stored at 4°C for further purification steps.

For deuteration in *E. coli,* we followed the protocol published by Sørensen *et al*.^20^ with slight variations. In brief, BL21 Star (DE3) cells transformed with the GbpA-encoding pET26-b(+) vector were grown for 5 h at 37°C, 220 rpm, in 2.5 mL LB medium containing 50 μg/mL Kan. 200 μL of this hydrogenated pre-culture was diluted in 2.5 mL of LB medium prepared with deuterium oxide as solvent (1/12.5 dilution) and grown for 6.5 h at 37°C, 220 rpm in presence of 50 μg/mL Kan. The deuterated pre-culture was then transferred to 25 mL (1/10 dilution) deuterated M9glyc+ minimal medium and grown for 14 h at 37°C, 130 rpm in presence of 50 μg/mL Kan. Deuterated M9_max_ media was produced following the recipe in Table S1, using anhydrous salts and deuterium oxide as solvent. Only the MEM vitamin solution and trace element solution contained water as solvent. The antibiotics solution was also prepared using deuterium oxide as solvent. The main culture was started by adding 225 mL of deuterated M9glyc+ medium to the culture (1/10 dilution) and adjusting the concentration of antibiotic back to 50 μg/mL Kan. IPTG was added to a final concentration of 1 mM to induce expression after 9.5 h, when the OD_600_ reached 2. After incubation for 20 h at 20°C, 130 rpm, cells were harvested and subjected to periplasmic extraction as described above for the hydrogenated protein, with all the centrifugation steps performed at 4,000 x *g* instead of 10,000 x *g*.

*Expression of cctx2*. C41 (DE3) cells (Sigma) transformed with pET24b(+) CCTX2 vector were grown overnight at 30°C, 120 rpm, in 50 mL LB medium^18^ with 100 μg/mL ampicillin (Amp). The main culture (1 L LB medium with 100 μg/mL Amp) was inoculated with the pre-culture to a 1/200 dilution factor and grown at 37°C, 110 rpm. When OD_600_ reached 0.6, the culture was cooled down in an ice bath for 10-15 min, and 0.5 mM IPTG was added to induce expression. After incubation for 20 h at 20°C, 110 rpm, cells were harvested at 4,500 x *g*, 4°C for 30 min. The cells were used immediately or stored at –80°C until use.

### Recombinant expression in Vibrio natriegens

#### Most cultures of Vmax™ X2 cells were grown in LB-v2 salts medium

*Expression of ctxAB and eltAB.* For CT and LT production, 10 mL of LB-v2 salts media containing 25 µg/mL CAM were inoculated with cells harboring the pARCT5 or pARpLT5 vectors and grown at 30°C and 180 rpm for 16 h. The cultures were diluted 1/100 in 500 mL of LB-v2 salts medium (25 µg/mL CAM) until OD_600_ reached ≈ 0.8 before induction with 0.2% L-arabinose (Sigma) at 30°C, 140 rpm for 20-22 h. CT and LT were harvested from the culture media by two rounds of centrifugation at 8,500 x g for 30 min at 4°C.

*Expression of gbpA*. GbpA was produced in minimal medium adapted for the growth of Vmax™ X2 (which we refer to as M9_max_) in baffled flasks (for media recipe, see Table S2). All media contained 200 μg/mL Kan, and incubation was carried out at 30°C and 120 rpm. Briefly, a pre-culture in 2 mL of LB-v2 salts medium was started from a glycerol stock and incubated for 5 h. A growth culture of 10 mL of minimal medium with 200 μg/mL Kan was then inoculated with 100 μL of the pre-culture (1/100 dilution), allowing it to grow for 14 h before topping it up with 90 mL of minimal medium. After 3h, *gbpA* expression was induced with the addition of 1 mM IPTG for 22 h, and the protein was harvested from the culture media by two rounds of centrifugation at 8,500 x *g* for 30 min at 4°C.

For deuteration in *V. natriegens,* we used the same recombinant construct, and adapted the protocol described above. All media contained 200 μg/mL Kan, and incubation was carried out at 30°C. Briefly, a pre-culture in 1 mL of LB-v2 salts medium was started from a glycerol stock and incubated for 3 h. Thereafter, 200 μL of this hydrogenated pre-culture was diluted in 2 mL of LB-v2 salts medium prepared with deuterium oxide as solvent (1/10 dilution) and grown for 4 h at 37°C, 220 rpm. The deuterated pre-culture was subsequently transferred to 10 mL (1/5 dilution) deuterated M9_max_ minimal medium and grown for 14 h at 110 rpm. Deuterated M9_max_ minimal medium was produced following the recipe in Table S2, using anhydrous salts and deuterium oxide as solvent. Only the MEM vitamin solution and trace element solution contained water as a solvent; the solution containing the antibiotic was prepared using deuterium oxide as solvent. The main culture was started by adding 90 mL of deuterated M9_max_ medium to the culture (1/10 dilution). IPTG was added to a final concentration of 1 mM to induce expression after 6 h. After incubation for 20 h at 120 rpm and 30°C. The culture media was harvested as described above by two rounds of centrifugation at 8,500 x *g* for 30 min at 4°C.

*Expression of cctx2*. For CCTX2, Vmax™ X2 cells transformed with pET24b(+) containing *cctx2* wild-type were grown at 30°C, 120 rpm overnight in presence of 400 μg/mL Kan. 1 L of LB-v2 salts medium (400 μg/mL Kan) was inoculated with 1/200 pre-culture and incubated at 30°C, 110 rpm. When the culture reached an OD_600_ of 0.8, expression was induced with 0.5 mM IPTG and were incubated at 30°C, 110 rpm for 18 h. Because CCTX2 is not secreted, cells were harvested at 8,500 x *g* for 30 min at 4°C, and the pellet was stored at –80°C until use.

### Protein purification

*Purification of CT and LT*. All steps were carried out at 4°C. CT was captured from the periplasmic fraction^22^ (*E. coli* expression) or the growth medium (Vmax™ X2 expression) by immobilized metal affinity chromatography (IMAC), as the toxin carries two histidine residues that confer natural weak affinity for Ni^2+^ and Co^2+^.^23^ The filtered solutions were directly applied onto a HiTrap Talon® crude 5 mL column (Cytiva) previously equilibrated with Talon A buffer, followed by a 15-column-volume (CV) wash with the same buffer and eluted with 10 CV Talon B (50 mM sodium phosphate pH 8.0, 300 mM NaCl, 50 mM imidazole). The protein was concentrated by ultrafiltration (4°C, 3,500 x *g*) using Amicon Ultra Centrifugal Filter Units 10K molecular weight cut-off (MWCO) (Merck). CT produced in *E. coli* was subsequently purified by cation-exchange chromatography with a HiTrap™ SP (GE Healthcare) column, followed by size-exclusion chromatography (SEC) with a Superdex 200 16/60 GL or Superdex 200 increase 10/300 GL column (Cytiva) equilibrated with Phosphate-Buffered Saline, pH 7.4 (PBS). For CT produced in Vmax™ X2, the cation exchange step was not needed. Fractions containing pure protein were pooled, concentrated by ultrafiltration as described above, and stored at 4°C.

As an alternative to IMAC, CT was captured from the medium by galactose affinity chromatography, exploiting the protein’s affinity for sugars. This was the primary method to capture LT from the medium, as it does not contain the histidine residues that confer affinity for divalent metals. Briefly, the filtered supernatant was loaded onto 3-4 mL immobilized D-galactose gel (Pierce™, Thermo Scientific™) equilibrated by gravity flow with Gal A buffer (50 mM Na-phosphate pH 7.4, 200 mM NaCl). After a 15 CV washing step with the same buffer, the protein was eluted with 10 CV of Gal B buffer (50 mM Na-phosphate pH 7.4, 200 mM NaCl, 300 mM D-galactose), concentrated by ultrafiltration and further purified by SEC in the same way as CT.

*Purification of GbpA.* To capture GbpA from the culture medium of Vmax™ X2, the medium was first dialyzed overnight at 4°C against 20 mM Tris-HCl pH 8.0, 100 mM NaCl (volume ratio 1:20 sample:buffer) using 10K MWCO SnakeSkin dialysis tubing (ThermoScientific). Dialyzed supernatant was subsequently loaded onto an equilibrated 5 mL HiTrap® Q XL column (Cytiva) for anion-exchange chromatography (AEX). For the protein produced in *E. coli*, the fractions resulting from the osmotic shock were directly loaded onto the AEX column. After a washing step with 20 CV binding buffer, the protein was eluted over a 12 CV 0-100% linear gradient with elution buffer (20 mM Tris-HCl pH 8.0, 400 mM NaCl). Fractions were analyzed by SDS-PAGE and those containing the target protein were pooled and concentrated with Amicon Ultra Centrifugal Filter Units 10K MWCO (Merck). GbpA was further purified by SEC using a Superdex 200 Increase 30/100 GL column (Cytiva) equilibrated with 20 mM Tris-HCl pH 8.0, 100 mM NaCl. For deuterated GbpA, a Superdex 75 Increase 30/100 GL column (Cytiva) equilibrated in the same buffer was used instead.

*Purification of CCTX2*. CCTX2 is retained in the cytoplasm of the cells. Cell pellets were resuspended in 5 mL lysis buffer (50 mM Na-HEPES, 200 mM NaCl, 2 mM EDTA, pH 7.5 supplemented with 5 mM dithiothreitol (DTT) and 1 x c*O*mplete protease inhibitor cocktail (Roche)) per gram of wet cell paste and incubated for 1 h at 4°C under stirring. The suspension was sonicated for 1 min (20% amplitude, 3 seconds on/7 seconds off) for *E. coli* cells and 5 min (30% amplitude, 3 seconds on/7 seconds off) for *V. natriegens* cells. The lysate was clarified at 40,000 x *g*, 4°C for 30 min to remove cell debris. The resulting supernatant containing soluble CCTX2 was then filtered through a 0.22 μm PES membrane filter and loaded onto a 5 mL Tricorn column packed with immobilized D-galactose gel (Pierce™, Thermo Scientific™) previously equilibrated with 6 CV loading buffer (20 mM Na-HEPES, 500 mM NaCl, pH 7.5). After a wash with 6 CV loading buffer, CCTX2 was eluted with 10 CV elution buffer (20 mM Na-HEPES, 500 mM NaCl, 1 M D-galactose, pH 7.5). The fractions containing CCTX2 were pooled and concentrated with an Amicon Ultra Centrifugal Filter Units 30K MWCO (Merck). Finally, CCTX2 was purified by SEC using a Superdex 200 Increase 30/100 GL column equilibrated with 50 mM Na-HEPES, 150 mM NaCl, 100 mM D-galactose, pH 7.5. In the case of the *E. coli* expression system, the whole fraction was loaded, whereas only half of the sample from the Vmax™ X2 expression system was used.

### Characterization of proteins and functional analysis

*Crystallization of CT, X-ray data collection and refinement*. Purified CT was dialyzed using a Pur-A-Lyzer™ Midi 3,500 MWCO (Sigma) into buffer G (50 mM Tris pH 7.4, 200 mM NaCl, 1 mM EDTA, 3 mM NaN_3_). The protein was crystallized by the sitting-drop vapor-diffusion method. Crystallization experiments were set up on 2-Lens UVXPO plates (SwissCI) using a crystallization robot (Oryx 4, Douglas Instruments) at 20°C. Crystals grew from 2 μL drops containing CT (5.76 mg/mL) and reservoir solution (0.125 M magnesium acetate, 24% w/v PEG 3350 and 300 mM D-galactose in Buffer G), mixed in a 1:1 volume ratio. Crystals were harvested with a nylon loop and cryo-protected in reservoir solution complemented with glycerol to a final concentration of 20% v/v. Cryo-protected crystals were flash-cooled in liquid nitrogen and shipped in a cryo-cooled dewar to a synchrotron light source for data collection. Data were collected at ID30B, ESRF, Grenoble (France). Diffraction images were integrated and scaled using *autoPROC*^24^; integrated and scaled intensities were merged and truncated with *AIMLESS*^25^ from the *CCP4* software suite.^26^ The resolution cut-off was chosen based on the CC_1/2_, which as described by Karplus & Diederichs,^27^ could be as low as 0.1 and still provide structural information. The structure was solved by molecular replacement using a previous CT structure determined to 1.9 Å (PDB ID: 1S5E)^28^ as search model in *Phaser*^29^ (*CCP4* suite).^26^ Refinement was performed by alternating cycles of manual rebuilding using *Coot,*^30^ and maximum-likelihood refinement with *REFMAC5.*^31^ Occupancy refinement was carried out with *phenix.refine* from the *Phenix* software suite.^32^ Data collection and refinement statistics are summarized in Table S3.

*Toxicity assays.* CHO cells seeded to a 24-well plate and grown overnight to 80% confluency were incubated for 2 h in serum-free medium containing 10-fold serial dilutions of CT, either purchased from Sigma (catalog #227036) or produced in Vmax™ X2. Cells were lysed in sample diluent from the Arbor Assays cAMP Direct ELISA, with clarified supernatants applied directly to the ELISA plate and processed for cAMP levels according to the manufacturer’s instructions. The cAMP content present in unintoxicated control cells was background subtracted from cAMP values recorded for the toxin-treated cells. The cAMP responses were then standardized to the maximal signal obtained with the commercial toxin by dividing the value from each technical replicate by the average cAMP response generated from cells exposed to 100 ng/mL of Sigma CT.

## RESULTS AND DISCUSSION

### Production of recombinant cholera toxin and heat-labile enterotoxin

In our optimized protocol for production of CT in *E. coli*, the protein was purified from the bacterial periplasmic space by IMAC, followed by ion-exchange and size-exclusion chromatography (Figure 1A-C). This protocol yielded only 0.2 mg of pure protein per L culture media, which was insufficient for our planned structural studies without significant scale-up. One of the main issues with this protocol was the presence of excess CTB compared to the CT holotoxin, which proved difficult to separate by chromatography techniques (Figure 1A, C).

**Figure 1.**
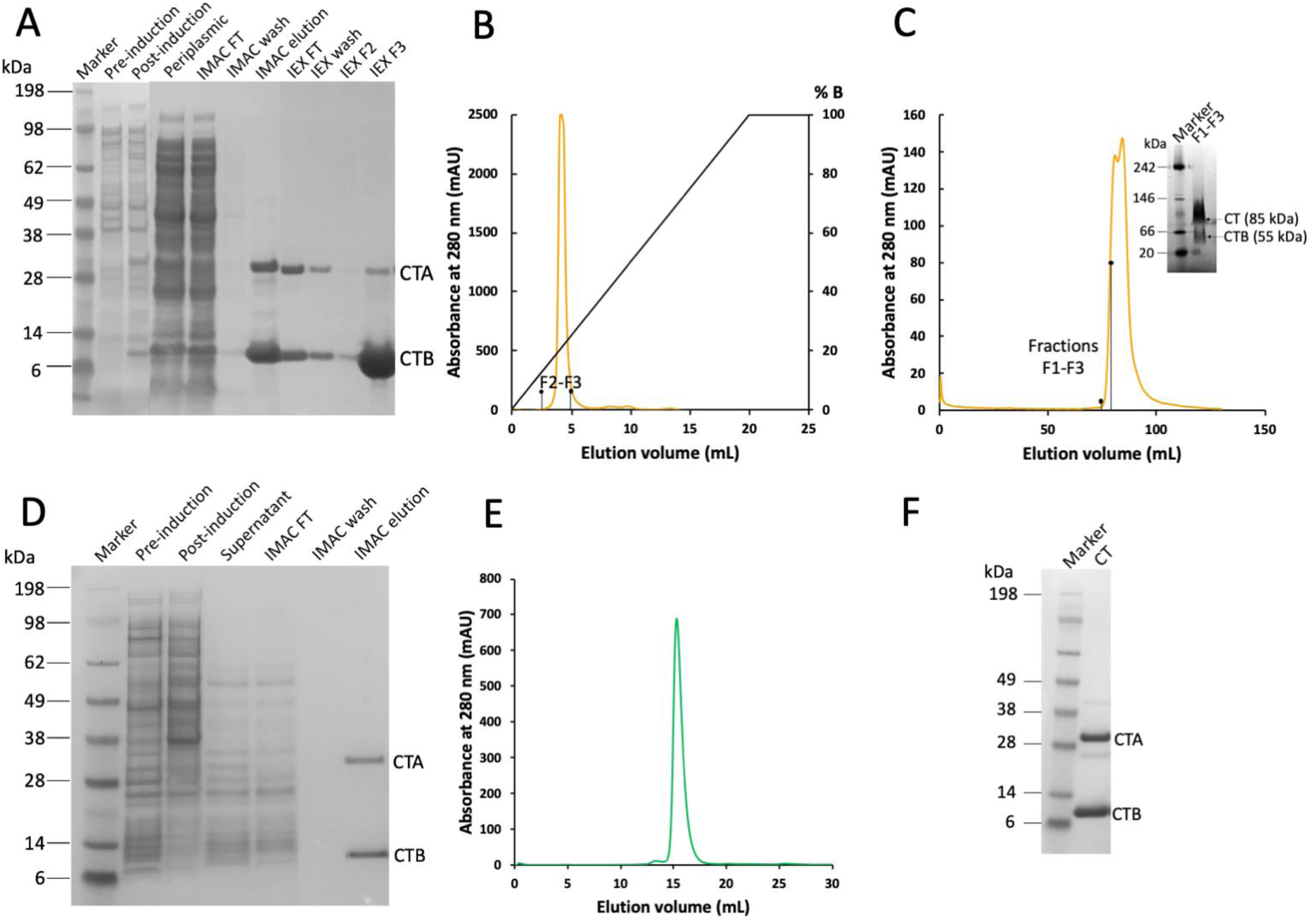
CT production in *E. coli* and Vmax™ X2. **A** SDS-PAGE analysis of CT samples obtained by expression in *E. coli* and purification by IMAC and ion-exchange chromatography. Periplasmic: periplasmic fraction, FT: flow-through. **B** Cation-exchange chromatogram and **C** SEC chromatogram of CT produced in *E. coli.* SEC fractions F1-F3 (first half of peak in C) on native PAGE (inset) still contained considerable amounts of CTB in the purified holotoxin sample. **D** SDS-PAGE gel from CT expression in Vmax™ X2, where CT is secreted into the growth medium. **E** SEC chromatogram of CT produced in Vmax™ X2, and **F** SDS-PAGE gel of SEC peak (compared to molecular mass marker).

In our efforts to increase CT yield and purity, we considered expressing the protein in a different host. Although toxigenic *V. cholerae* strains have been previously used to produce and purify CT,^33–37^ they are inherently pathogenic and are limited to production of the native toxin. It is of note that also non-toxigenic *V. cholerae* strains have been used for producing recombinant CTB,^38–40^ however, transforming such strains with the desired plasmids proved challenging.^38^ To the best of our knowledge, these strains have not been used for producing CT.

As an alternative, we decided to test expression in Vmax™ X2, an engineered strain derived from the non-pathogenic bacterium *Vibrio natriegens*^5^ (Figure 1D-F). Since this bacterium is a close relative of *V. cholerae*, we presumed that higher CT amounts could be obtained when using this host instead of *E. coli*. We also hypothesized that CT could be secreted to the extracellular environment when produced in Vmax™ X2, as it contains the T2SS used by *V. cholerae* to secrete CT.^14,41,42^ Although such a machinery also exists in *E. coli*, CT is not secreted in soluble form in this expression system,^43^ possibly because the toxins remain associated with the *E. coli* surface *via* binding to their outer-membrane lipopolysaccharides (LPS).^44,45^ In contrast, CT (and LT) do not have affinity for *Vibrio* LPS,^44^ which would explain why the toxin can be secreted as a soluble protein in *Vibrio* strains. Using the same plasmids that we previously used for *ctxAB* expression in *E. coli*, we developed a protocol for production of CT in *V. natriegens* that we also adapted for production of other proteins from the same species (bacterial protein GbpA; including protein deuteration), from another phylum (*E. coli* LT) or from another domain of life (the fungal toxin CCTX2).

As a halophilic bacterium, Vmax™ X2 requires high salt concentrations for optimal growth. Therefore, cells were grown in the common LB medium supplemented with additional “v2” salts (see *Materials and Methods*), as recommended by the manufacturer.^46^ Since Vmax™ X2 can produce proteins at a wide range of optical density values (OD_600_) with little variations in yield,^5,46^ we decided to induce expression at the standard OD_600_ value 0.8. Due to the fast growth of Vmax™ X2, such density was quickly achieved, 2 h after inoculating the main culture. In our initial test, expression was induced with 0.2% L-arabinose (the same concentration used for induction in *E. coli*), and samples before and after 20 h induction were collected and analyzed by SDS-PAGE. CT production was not visible ‒ neither in pre or post-induction cell samples, nor in the culture supernatant (Figure 1D, first three lanes after marker). This was not surprising, as we expected the protein to be secreted into the growth medium, and it may be too diluted to be clearly seen in the gel. To verify that CT was indeed produced, the supernatant was directly loaded onto an IMAC column, and the elution fraction was analyzed by SDS-PAGE. Two clear bands matching the sizes of CTA and CTB were observed (Figure 1D, last lane), confirming successful production of the toxin. CT appears to already be almost pure after this step; nevertheless, the protein was subjected to SEC to assess the presence of CTB and remove aggregates or contaminants that may not be visible on the gel. A single sharp peak was obtained (Figure 1E, with corresponding SDS-PAGE gel in Figure 1F), suggesting that there was little or no contamination by CTB, a major issue for expression in *E. coli* (Figure 1 C). It has been observed earlier that CT holotoxin (or ‘*choleragen’*) production simultaneously yields CTB (also referred to as ‘*choleragenoid’*).^34^ Subsequently, the subunit assembly process has been characterized, mainly based on studies of LT,^35,47,48^ and it was confirmed that B-pentamers can form and be exported on their own, whereas uncomplexed A-subunits remain associated with the cells.^35^ Interestingly, assembly into holotoxins was shown to be promoted by the A-subunit,^47,48^ which on its own is highly unstable and prone to degradation. Since expression in Vmax™ X2 was performed at a lower temperature compared to *E. coli* (30°C *versus* 37°C in *E. coli*), we suspect that CTA is less affected by thermal degradation in this system, leading to a more homogenous production of the holotoxins. Although the protein yields varied significantly between different expression rounds, at least 10 times more CT was obtained in Vmax™ X2 compared to *E. coli* (2 mg per L medium in Vmax™ X2 vs. 0.2 mg per L medium in *E. coli*). By inducing expression with 0.02% L-arabinose and loading the medium on a galactose resin instead of IMAC, up to 20% higher yields were obtained. This protocol could be further optimized, but in its current form it already allowed to obtain sufficient amounts of toxin for structural studies. Furthermore, secretion of the protein to the growth medium simplified the downstream purification procedure and made it more time-efficient.

To verify that the CT holotoxin produced in Vmax™ X2 was correctly folded, we determined its crystal structure to 2.3 Å resolution (Figure 2). The structure was refined to *R*/*R*_free_ values of 22.1/26.2%, and exhibited good geometry based on the Ramachandran plot as well as small deviations from ideal bond lengths and angles (Table S3). The crystal structure is essentially identical to the CT crystal structure published earlier by the Hol lab (Figure 2), from protein produced in *E. coli,*^28^ with an RMSD value for all C_α_ atoms of 0.3 Å, confirming the correct folding of the holotoxin.

**Figure 2.**
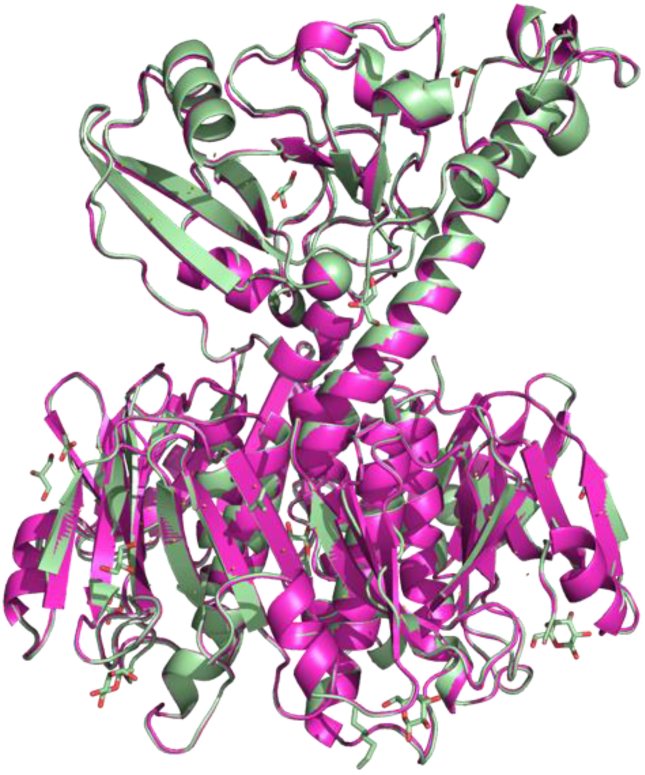
Comparison of CT crystal structures. CT produced in Vmax™ X2 (light green; PDB ID: 8QRE, this work), superimposed onto the published crystal structure of CT produced in *E. coli* (magenta; PDB ID: 1S5E^28^). RMSD: 0.3 Å.

We also confirmed that the produced toxin retains its biological activity. In fact, the toxin produced in our lab elicited a stronger cAMP response in intoxicated cells than CT obtained from a commercial supplier (Figure 3A). This difference likely relates to the greater quantity of intact holotoxin in the Vmax™ X2 preparation when compared to the commercial preparation: both preparations had equivalent levels of the CTB subunit as assessed by SDS-PAGE, but the commercial preparation had a lower quantity of CTA and, thus, less functional toxin (Figure 3B). Additionally, commercial CT is delivered as a lyophilized sample, increasing the risk of denaturation.

**Figure 3.**
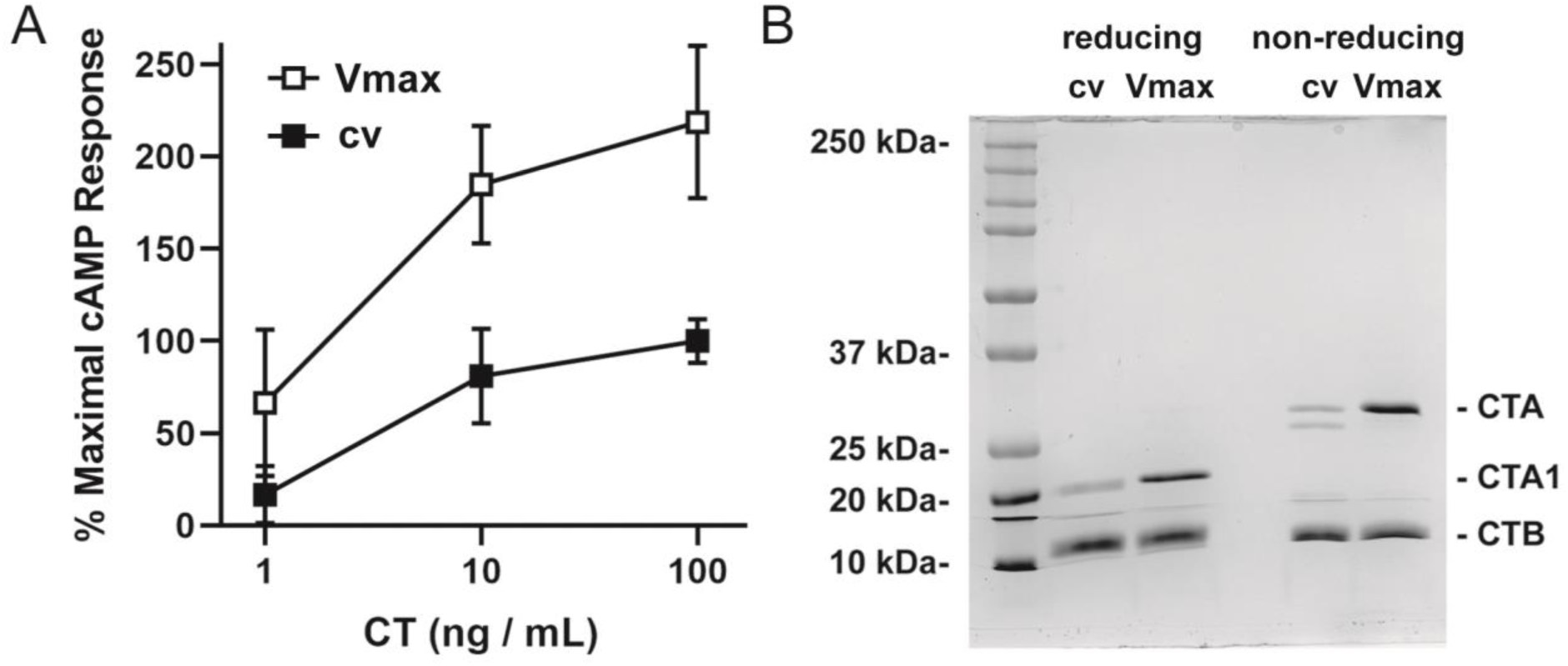
Toxicity and stoichiometry for commercial and Vmax™ X2 preparations of CT. **A** CHO cells were incubated for 2 h with 10-fold dilutions of CT purchased from a commercial vendor (cv, filled squares) or purified from Vmax™ X2 (Vmax, open squares). An ELISA was then used to quantify cAMP levels from the intoxicated cells. Background-subtracted data were expressed as percentages of the response elicited from cells challenged with 100 ng/mL of the commercial toxin and represent the means ± standard deviations of nine technical replicates from three independent experiments. **B** Samples of CT purchased from a commercial vendor (cv) or produced in Vmax™ X2 (Vmax) were resolved by SDS-PAGE under reducing and non-reducing conditions. The samples (4 μg per lane) were visualized by Coomassie stain. The entire gel is shown, with a listing of the molecular masses from select protein standards.

A close homolog of CT is LT from ETEC.^49^ Although we had not produced LT in *E. coli* before, our promising results using Vmax™ X2 for expression of *ctxAB* encouraged us to directly test this host for *eltAB* expression; specifically, we produced pLT, the toxin originating from ETEC strains infecting pigs. Since CT and LT share more than 80% sequence identity,^49^ we applied the same Vmax™ X2 expression protocol developed for CT. Like CT, pLT was secreted into the medium and, after purification by galactose affinity chromatography (Figure 4), comparable yields to CT were obtained (7 mg of protein per L medium).

**Figure 4.**
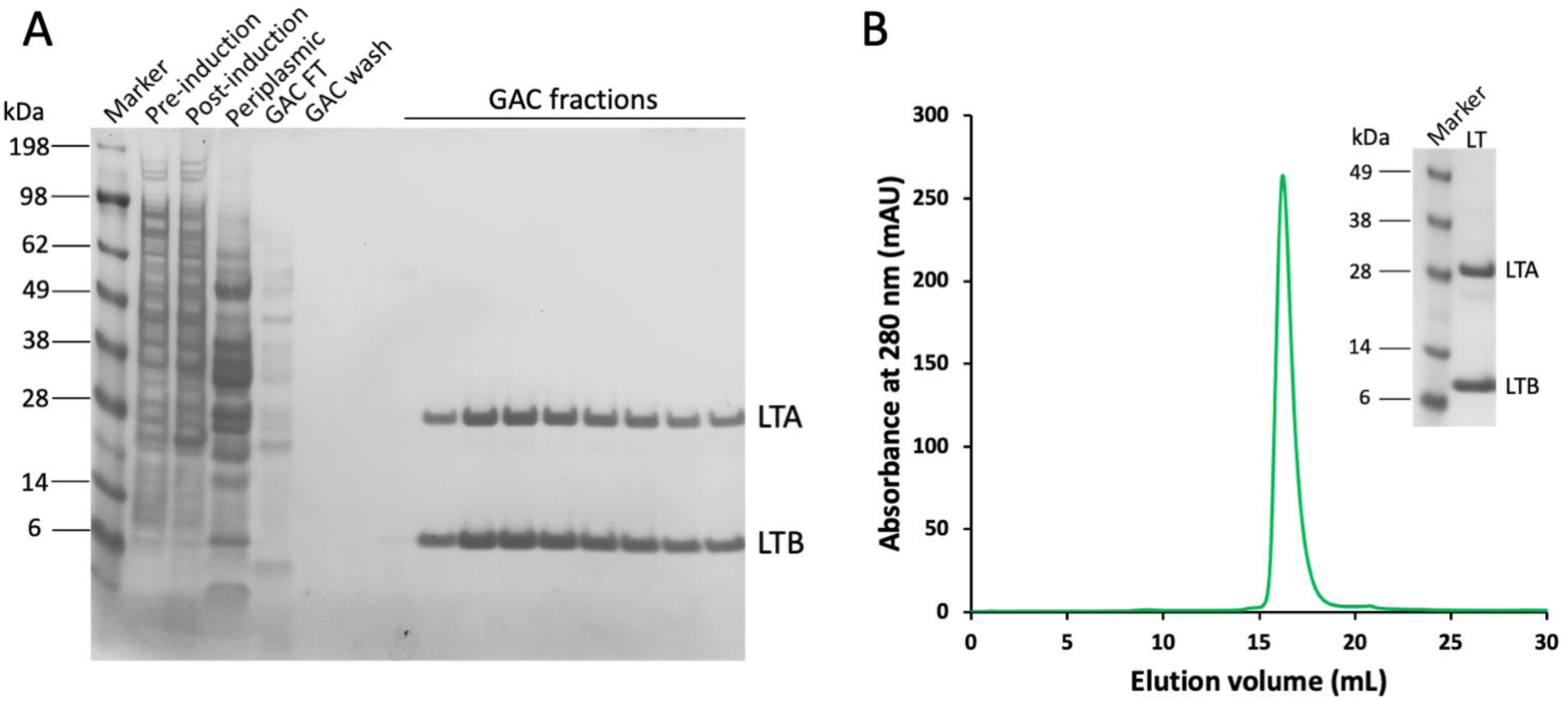
pLT production in Vmax™ X2. **A** SDS-PAGE analysis of pLT samples obtained for expression in Vmax™ X2 and purification from the culture supernatant by galactose-affinity chromatography (GAC). **B** SEC chromatogram of pLT captured by GAC, and SDS-PAGE of SEC fractions (inset).

### Production of another V. cholerae protein: GbpA

Another protein that is natively secreted by the T2SS is the *V. cholerae* colonization factor GbpA. This adhesin binds to chitin and mucins, helping the pathogen to survive in its natural marine environment and to colonize the human intestine, respectively.^50^ In our lab, GbpA has routinely been produced in *E. coli*, in both TB and minimal media,^20^ and purified from the *E. coli* periplasmic fraction by ion-exchange chromatography and SEC. Interestingly, we found in previous work that better yields were obtained when producing the protein in minimal media instead of the nutrient-rich media TB (15 mg per L in M9 vs 7 mg per L in TB).^20^ We suspected that this might be due to misfolding of GbpA when produced too quickly in TB, whereas slower expression in M9 would result in less aggregates. Although GbpA was already produced in relatively high yields in *E. coli*, we decided to test expression in Vmax™ X2, given our encouraging results with the bacterial toxins (Figures 2 and 4). When expressed in Vmax™ X2, GbpA could be seen in the culture supernatant (Figure S2), indicating that the protein is recognized by the T2SS machinery of *V. natriegens* and secreted. GbpA is produced in much greater amounts than CT, which is not surprising since *gbpA* is also highly expressed in *E. coli*. Production in Vmax™ X2 not only increased GbpA yield by more than 6-fold compared to production in *E. coli*, but also simplified protein purification. SEC chromatograms and SDS-PAGE analysis of the produced samples are shown in Figure 5A-C. Similar results were recently reported in an independent study by Schwarz *et al*., who used GbpA as a secretion and affinity purification tag for an antimicrobial peptide produced in *V. natriegens.*^11^ Because GbpA is isolated from the culture supernatant, contaminants from the periplasm are avoided, and aggregation is less likely to occur. These results demonstrate that Vmax™ X2 is a suitable host to express virulence factors secreted by the T2SS, both from *V. cholerae* and other bacteria like ETEC.

**Figure 5.**
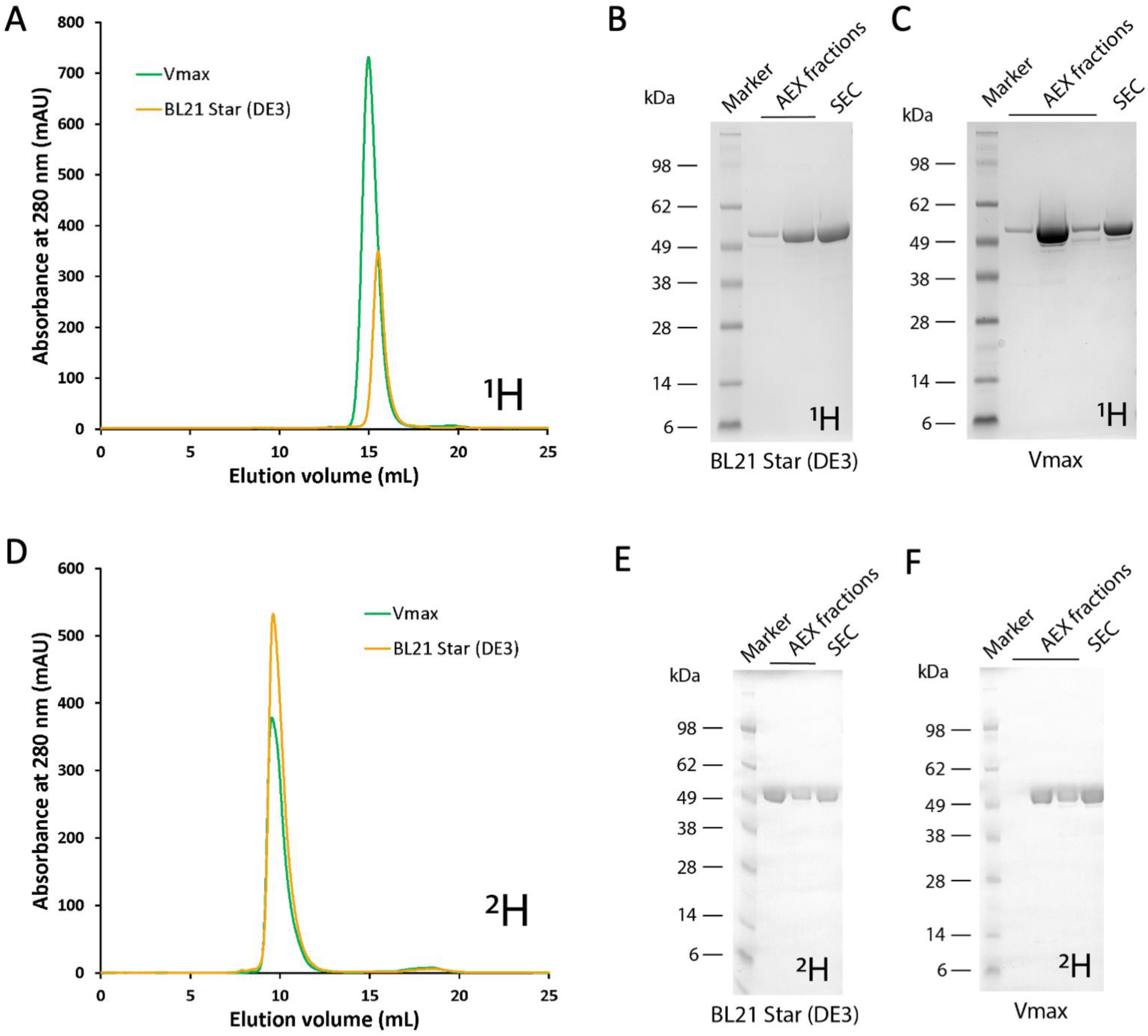
GbpA production in *E. coli* and Vmax™ X2. **A** SEC chromatogram (Superdex 200 Increase 30/100 GL column) of hydrogenated (^1^H) GbpA expressed in *E. coli* BL21 Star(DE3) (light orange) and in Vmax™ X2 (green). Several SEC runs were performed for each purification batch; therefore the intensity of absorbance is not correlated with yield. **B-C** SDS-PAGE analysis of GbpA hydrogenated (^1^H) samples expressed in Vmax™ X2 or *E. coli* BL21 Star(DE3) after the AEX and SEC purification steps. **D** SEC chromatogram (Superdex 75 Increase 30/100 GL column) of deuterated (^2^H) GbpA expressed in *E. coli* BL21 Star(DE3) (light orange) and in Vmax™ X2 (green). Again, SEC was performed in several injections per batch and the amounts shown do not correlate with yield. **E-F** SDS-PAGE analysis of GbpA deuterated (^2^H) samples expressed in Vmax™ X2 or *E. coli* BL21 Star(DE3) after the AEX and SEC purification steps. ^1^H refers to ‘common’ hydrogen (also called protium); ^2^H to deuterium.

Since we intend to perform neutron scattering experiments of GbpA,^20^ we would strongly benefit from producing perdeuterated protein. However, this is not necessarily easy to achieve, and also quite expensive. Given our encouraging results using *V. natriegens* for producing hydrogenated GbpA, we therefore tested *gbpA* expression with the Vmax™ X2 system, using deuterated minimal media and deuterium oxide as solvent. Indeed, we were able to produce deuterated GbpA using Vmax™ X2 in almost 3-fold higher yields than in *E. coli* (41.6 mg per L in Vmax™ X2 vs. 14.8 mg per L in *E. coli*). Moreover, downstream processing of the produced protein was simplified, as was the case for hydrogenated media. SEC chromatograms and SDS-PAGE analysis of the deuterated samples are shown in Figure 5D-F.

### Production of a non-secreted fungal protein toxin: CCTX2

To test the full potential of the Vmax™ X2 system, we applied our protocol to purify CCTX2, a fungal nematotoxin, for which no structural information is available to date, and very little is known about its function. Preliminary work in both our own lab at UiO and that of our collaborators at ETH Zurich (Künzler lab) had yielded only low amounts of CCTX2 in *E. coli* (approximately 0.4 mg per 1 L culture media). Since structural characterization often requires large amounts of protein, this seemed to be a perfect test case. We carried out *cctx2* expression in Vmax™ X2 and compared its yield in post-induction samples to *E. coli* C41(DE3), which was previously selected as the best-expressing strain from a large library of *E. coli* strains. As seen in Figure 6, Vmax™ X2 cells clearly produced a greater amount of protein than the *E. coli* system. Purification was performed using the same protocol for both strains, revealing the greater potential of Vmax™ X2 already at the stage of galactose affinity chromatography. The elution profile shows a very high amount of protein, with the UV_280_-detector reaching saturation for protein expressed in Vmax™ X2 (Figure 6A). Purification by SEC on Superdex 200 Increase 30/100 GL resulted in large amounts of pure CCTX2, with higher A_280_ absorption intensity compared to C41(DE3) cells (Figure 6B). This was confirmed by SDS-PAGE (Figure 6C,D). After SEC, the sample from Vmax™ X2 provided consistent yields of approximately 10 mg of protein per liter culture. The new protocol thus resulted in a 26-fold increase in yield.

**Figure 6.**
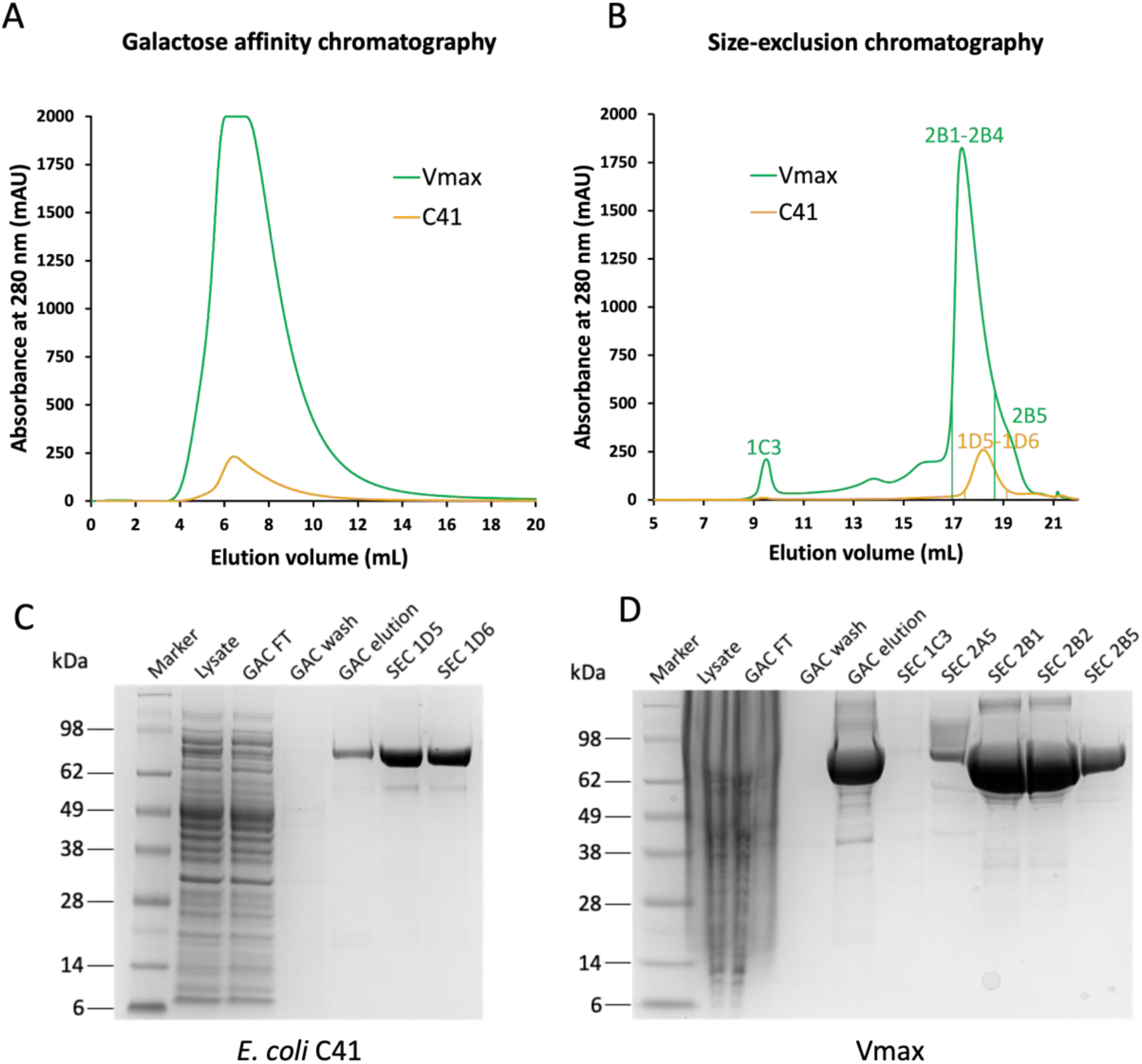
CCTX2 production in *E. coli* and Vmax™ X2. **A** Galactose-affinity chromatography of CCTX2 produced in 1 L cultures from *E. coli* (light orange) and *V. natriegens* (green). **B** Matching SEC chromatogram comparing production in *E. coli* (light orange) and *V. natriegens* (green) for 0.5 L cultures. **C-D** SDS-PAGE of CCTX2 samples from expression in *E. coli* C41(DE3) and Vmax™ X2, matching purification fractions labeled in panel B.

## CONCLUSION

*V. natriegens* proved to be a highly advantageous host for high-yield production of virulence factors naturally secreted by the T2SS. In Vmax™ X2, the yield of CT was 10-fold higher when compared to *E. coli*, and a 6-fold increase was obtained for GbpA (3-fold increase for GbpA deuteration). Moreover, secretion of the recombinant proteins simplified the purification process and, in several cases, improved sample purity, since less contaminants are present in the culture supernatant compared to the bacterial cytosol and periplasm. For the fungal toxin CCTX2, a test case representing non-secreted proteins, Vmax™ X2 even allowed us to increase the production efficiency by 26-fold, which greatly benefits our ongoing structural characterization. We hope that our work encourages the use of *V. natriegens* as an alternative expression host for the production of other difficult expression targets, not only from *Vibrio* species but also from other bacteria and fungi. *V. natriegens* is also useful for protein deuteration and for isotope labeling in general.

## SUPPORTING INFORMATION

This article contains supporting information (Supporting Figures S1 to S2 and Supporting Tables S1 to S3).

*Figure S1*. Schematic plasmid maps showing some major features

*Figure S2.* SDS-PAGE analysis of GbpA fractions before purification

*Table S1*. Minimal medium M9glyc+ for GbpA production in BL21(DE3)

*Table S2*. Minimal medium M9max for GbpA production in Vmax™ X2

*Table S3*. Data collection and refinement statistics for CT produced from Vmax™ X2

### Accession IDs

UniProt IDs: P01555 and E9RIX3 (Genes *ctxA* and *ctxB*). PDB ID: xxxx

UniProt IDs: P06717 and P32890 (Genes *eltA* and *eltB*)

UniProt ID: Q9KLD5 (Gene *gbpA*).

UniProt ID: A8NDT7 (Locus tag CC1G_10077, gene *cctx2*).

### ABBREVIATIONS

AEX, anion-exchange chromatography; CT, cholera toxin; CTA, catalytic A-subunit of CT; CTB, homopentamer of receptor-binding B-subunits of CT; CCTX2, *Coprinopsis cinerea* toxin 2; EDTA, ethylenediaminetetraacetic acid; ESRF, European Synchrotron Radiation Facility; GAC, galactose-affinity chromatography; GbpA, *N*-acetylglucosamine-binding protein A; HEPES, N-(2-hydroxyethyl)piperazine-N′-(2-ethanesulfonic acid); IMAC, ion-immobilized affinity chromatography; IPTG, isopropyl β-D-1-thiogalactopyranoside; LB, Luria Bertani; MWCO, molecular weight cut-off; OD, optical density; PBS, Phosphate-Buffered Saline pH 7.4; PMSF, phenylmethylsulfonyl fluoride; PES, polyethersulfone; SDS-PAGE, sodium dodecyl sulfate– polyacrylamide gel electrophoresis; SEC, size-exclusion chromatography; TB, terrific broth; Tris, tris(hydroxymethyl)aminomethane; TT2S, Type II secretion system; Vmax, *Vibrio natriegens*

## AUTHOR CONTRIBUTIONS

U.K. conceived the study, and N.M. piloted the work. N.M., F.K. and M.M.-C. performed most of the experimental work (expression in *E. coli* and Vmax, and purification; M.M.-C. performed protein deuteration), supervised by U.K.. A.M.T. performed initial experiments for GbpA during her Master’s project, supervised by U.K.. N.M. determined the crystal structure of CT, supervised by G.C. and U.K., who also validated the structure. G.R.H.III performed the toxicity assays, supervised by K.T. The first draft of the manuscript was written by N.M., and revised by N.M, F.K., M.M.-C, G.C, K.T. and U.K., with final approval by all authors.

## ACKNOWLEDGEMENTS

We thank Drs. Randall K. Holmes (University of Colorado School of Medicine) and Markus Künzler (ETH Zurich) for kindly sharing the plasmids for expression of *ctxAB* and *cctx2*, respectively. In addition, we thank previous members of our group, especially Henrik V. Sørensen and Joel B. Heim, for their contributions as co-supervisors of Master’s students and for comments on the manuscript. X-ray diffraction experiments were performed on beamline ID30B2 at the European Synchrotron Radiation Facility (ESRF), Grenoble, France. We are grateful to Local Contact Christoph Mueller-Dieckmann at the ESRF, and Tamjidmaa Khatanbataar, for providing assistance in using the beamlines and data collection. Work at UiO was performed at the UiO Structural Biology core facilities.

## FUNDING INFORMATION

This work was supported by the National Institutes of Health (award number R01AI137056 to our collaborator Dr. Ken Teter), by the Norwegian Research Council (grant no. 272201 to UK) and by the University of Oslo (PhD positions of N.M., F.K. and M.M.-C.). The content is solely the responsibility of the authors and does not necessarily represent the official views of the funding agencies. Most of the work was carried out at the UiO Structural Biology core facilities, which are part of the Norwegian Macromolecular Crystallography Consortium NORCRYST and received funding from the Norwegian INFRASTRUKTUR-program (project no. 245828) as well as from UiO (core facility funds).

## CONFLICT OF INTEREST

The authors declare that they have no conflicts of interest with the contents of this article.

## SUPPORTING INFORMATION

**Figure S1.**
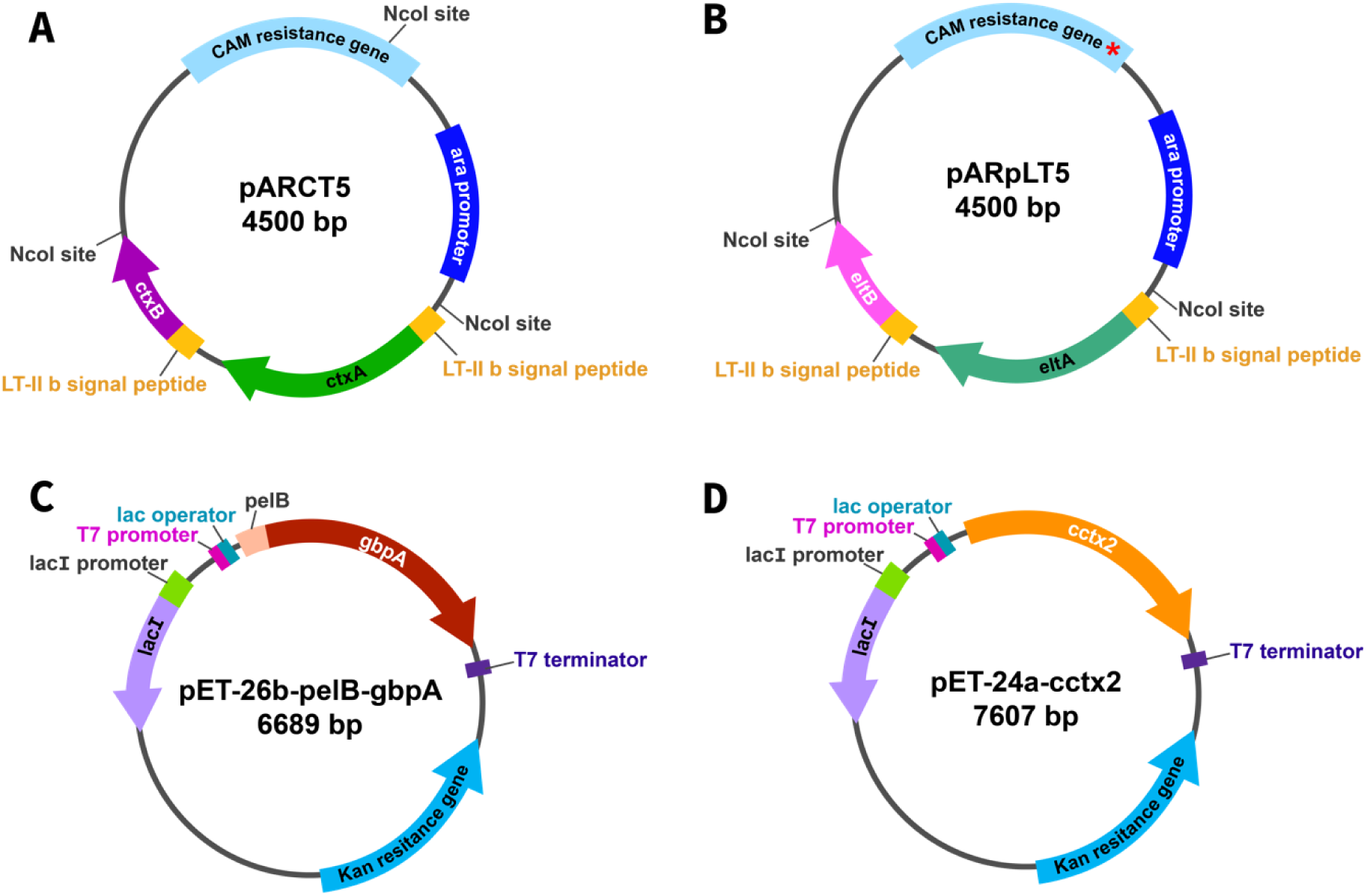
Plasmid maps. **A** *ctxAB* expression plasmid (pARCT5). **B** *eltAB* expression plasmid (pARpLT5). The red asterisk in the CAM resistance gene represents the silent mutation introduced to remove an NcoI site (leaving only two NcoI restriction sites in the vector). **C** *gbpA* expression plasmid (pET-26b-pelB-gbpA). **D** *cctx2* expression plasmid (pET-24-a-cctx2)

**Figure S2.**
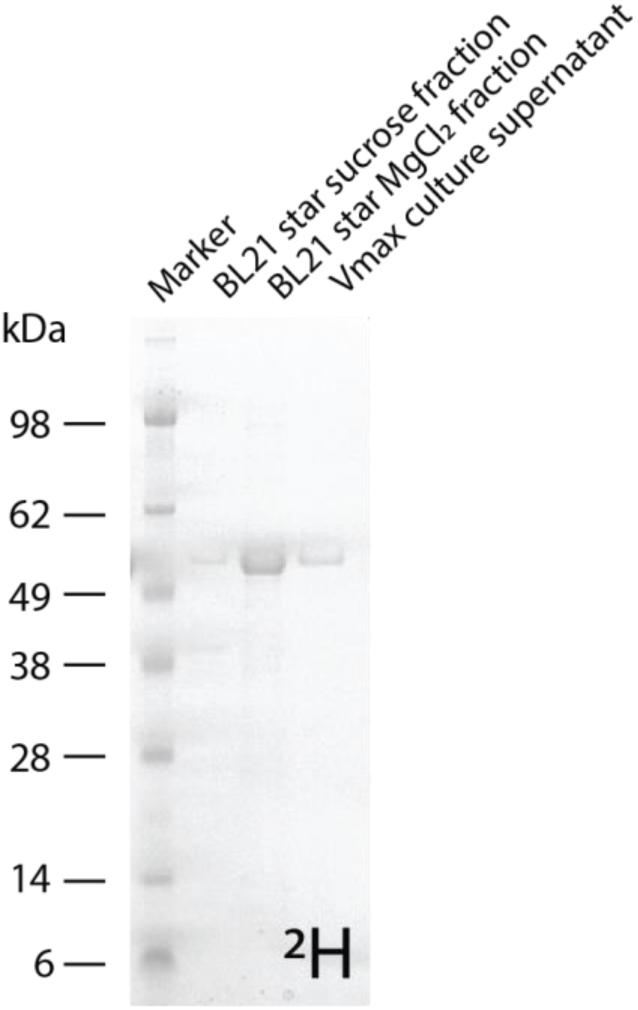
SDS-PAGE analysis of GbpA fractions before purification. ^2^H is the chemical symbol for deuterium, a chemical isotope of hydrogen with one proton and one neutron (whereas normal hydrogen (^1^H) lacks neutrons).

**Table S1:** M9glyc+ minimal medium for GbpA production in BL21(DE3)

| Minimal medium |  |
| --- | --- |
| K<sub>2</sub>HPO<sub>4</sub> | 109 mM |
| KH<sub>2</sub>PO<sub>4</sub> | 37 mM |
| Na<sub>2</sub>HPO<sub>4</sub> | 63 mM |
| K<sub>2</sub>SO<sub>4</sub> | 14 mM |
| NH<sub>4</sub>Cl | 93 mM |
| MgCl<sub>2</sub><sup>a</sup> | 10 mM |
| Glycerol | 1.6 % |
| MEM vitamins<sup>b</sup> | 1x |
| Trace Element solution<sup>c</sup> | 1x |

<sup>a</sup> Added from a stock solution just before inoculation to minimize precipitation of phosphate salts
<sup>b</sup> MEM 100x vitamin solution from Sigma-Aldrich
<sup>c</sup> The 1000x trace element solution was made by dissolving the following ingredients in 100 mL H<sub>2</sub>O: 0.6 g FeSO<sub>4</sub>×7H<sub>2</sub>O, 0.6 g CaCl<sub>2</sub>×2H<sub>2</sub>O, 0.12 g MnCl<sub>2</sub>×4H<sub>2</sub>O, 0.08 g CoCl<sub>2</sub>×6H<sub>2</sub>O, 0.07 g ZnSO<sub>4</sub>×7H<sub>2</sub>O, 0.03 g CuCl<sub>2</sub>×2H<sub>2</sub>O, 2 mg H<sub>3</sub>BO<sub>4</sub>, 0.025 g (NH<sub>4</sub>)<sub>6</sub>Mo<sub>7</sub>O<sub>24</sub>×4H<sub>2</sub>O, 0.5 g EDTA.

**Table S2:**
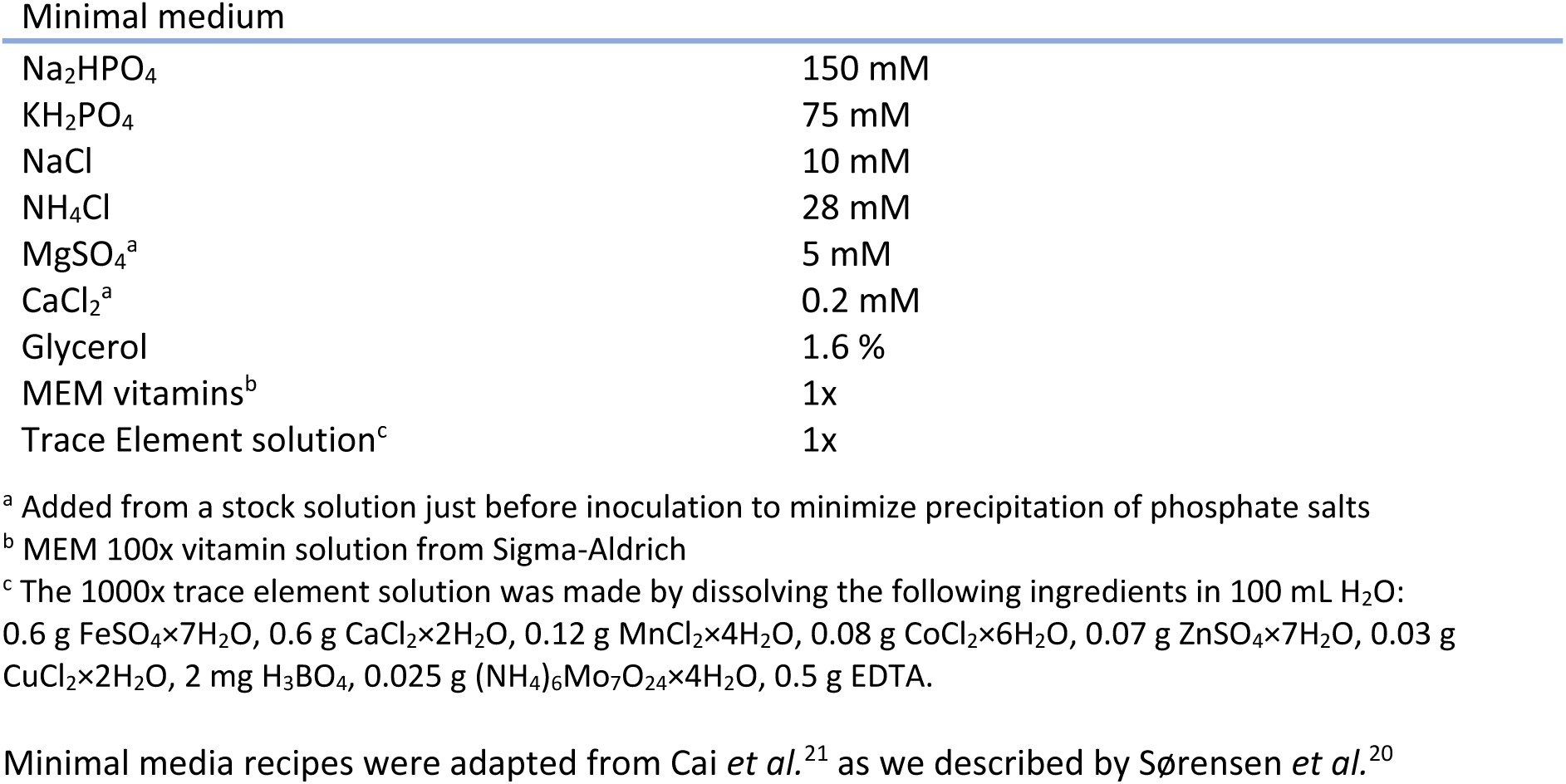
M9_max_ minimal medium for GbpA production in Vmax™ X2.

| Minimal medium |  |
| --- | --- |
| Na<sub>2</sub>HPO<sub>4</sub> | 150 mM |
| KH<sub>2</sub>PO<sub>4</sub> | 75 mM |
| NaCl | 10 mM |
| NH<sub>4</sub>Cl | 28 mM |
| MgSO<sub>4</sub><sup>a</sup> | 5 mM |
| CaCl<sub>2</sub><sup>a</sup> | 0.2 mM |
| Glycerol | 1.6 % |
| MEM vitamins<sup>b</sup> | 1x |
| Trace Element solution<sup>c</sup> | 1x |

<sup>a</sup> Added from a stock solution just before inoculation to minimize precipitation of phosphate salts
<sup>b</sup> MEM 100x vitamin solution from Sigma-Aldrich
<sup>c</sup> The 1000x trace element solution was made by dissolving the following ingredients in 100 mL H<sub>2</sub>O: 0.6 g FeSO<sub>4</sub>×7H<sub>2</sub>O, 0.6 g CaCl<sub>2</sub>×2H<sub>2</sub>O, 0.12 g MnCl<sub>2</sub>×4H<sub>2</sub>O, 0.08 g CoCl<sub>2</sub>×6H<sub>2</sub>O, 0.07 g ZnSO<sub>4</sub>×7H<sub>2</sub>O, 0.03 g CuCl<sub>2</sub>×2H<sub>2</sub>O, 2 mg H<sub>3</sub>BO<sub>4</sub>, 0.025 g (NH<sub>4</sub>)<sub>6</sub>Mo<sub>7</sub>O<sub>24</sub>×4H<sub>2</sub>O, 0.5 g EDTA.

**Table S3.** Data collection and refinement statistics for CT produced in Vmax™ X2.

| **Data collection** |  |
| --- | --- |
| Beamline | ESRF-ID30B |
| Wavelength (Å) | 0.9763 |
| Space group | $P2_1$ |
| Cell parameters – a, b, c (Å) | 60.7, 108.1, 124.8 |
| Protein chains in a.u. | 2 |
| Matthew's coefficient (Å<sup>3</sup>/Da) | 2.4 |
| Resolution (Å)<sup>a</sup> | 124.1-2.3 (2.35-2.30) |
| CC<sub>1/2</sub> (%)<sup>a</sup> | 97.3 (17.0) |
| $R_{\text{merge}}$ (%)<sup>a</sup> | 25.6 (>100) |
| $R_{\text{p.i.m}}$ (%)<sup>a</sup> | 16.9 (>100) |
| Mean $I/\sigma(I)$<sup>a</sup> | 4.3 (0.6) |
| Completeness (%)<sup>a</sup> | 98.6 (99.3) |
| Number of unique reflections<sup>a</sup> | 70156 (4548) |
| Multiplicity<sup>a</sup> | 3.0 (2.9) |
| Wilson $B$-factor (Å<sup>2</sup>) | 38.7 |
| **Refinement** |  |
| $R_{\text{work}}/R_{\text{free}}$ (%) | 22.1/26.2 % |
| Average $B$-factor (Å<sup>2</sup>) | 44.7 |
| Number of atoms | 13002 |
| Protein | 12409 |
| Water | 354 |
| Ligands | 239 |
| r.m.s.d from ideal geometry |  |
| Bond lengths (Å) | 0.01 |
| Bond angles (deg.) | 1.14 |
| Ramachandran plot |  |
| Favored (%) | 96.9 |
| Allowed (%) | 3.0 |
| Outliers (%) | 0.1 |
| PDB ID | 8QRE |

<sup>a</sup> Values in parenthesis refer to the highest-resolution shell

## Notes

### Competing Interest Statement

The authors have declared no competing interest.

